# The dynamic surface proteomes of allergenic fungal conidia

**DOI:** 10.1101/2020.01.10.902015

**Authors:** Matthew G. Blango, Annica Pschibul, Flora Rivieccio, Thomas Krüger, Muhammad Rafiq, Lei-Jie Jia, Tingting Zheng, Marie Goldmann, Vera Voltersen, Jun Li, Gianni Panagiotou, Olaf Kniemeyer, Axel A. Brakhage

## Abstract

Fungal spores and hyphal fragments play an important role as allergens in respiratory diseases. In this study, we performed trypsin shaving and secretome analyses to identify the surface-exposed proteins and secreted/shed proteins of *Aspergillus fumigatus* conidia, respectively. We investigated the surface proteome under different conditions, including temperature variation and germination. We found that the surface proteome of resting *A. fumigatus* conidia is not static, but instead unexpectedly dynamic, as evidenced by drastically different surface proteomes under different growth conditions. Knockouts of two abundant *A. fumigatus* surface proteins, ScwA and CweA, were found to function only in fine-tuning the cell wall stress response, implying that the conidial surface is very robust against perturbations. We then compared the surface proteome of *A. fumigatus* to other allergy-inducing molds, including *Alternaria alternata, Penicillium rubens*, and *Cladosporium herbarum*, and performed comparative proteomics on resting and swollen conidia, as well as secreted proteins from germinating conidia. We detected 125 protein ortholog groups, including 80 with putative catalytic activity, in the extracellular region of all four molds, and 42 nonorthologous proteins produced solely by *A. fumigatus*. Ultimately, this study highlights the dynamic nature of the *A. fumigatus* conidial surface and provides targets for future diagnostics and immunotherapy.

## Introduction

120,000 fungal species have been described to date; however, only a few hundred of these fungi are able to cause illness in humans.^1^ These diseases range from mild allergic reactions to life-threatening systemic infection. In terms of allergy, more than 80 mold genera have been associated with hypersensitive responses in allergic patients.^2^ Within this group, species of the genera *Alternaria* and *Cladosporium* represent the most important outdoor allergens, while species of *Penicillium* and *Aspergillus* are reported to be the most common indoor airborne fungi.^3^ *A. fumigatus* is associated with different forms of allergic diseases, including allergic bronchopulmonary aspergillosis (ABPA;^4^), a persistent colonization of the bronchi known to frequently occur in patients with asthma and cystic fibrosis (CF). In addition, *A. fumigatus* is the most common species within the mold fungi to cause invasive infections in the lungs of immunocompromised patients, which is designated invasive aspergillosis.^5–7^

The *A. fumigatus* infectious lifecycle starts with the inhalation of conidia, 2-3 μm asexual spores, produced by mature conidiophores.^8^ After deposition in the lungs, conidia undergo germination, whereby they initially swell, extend germ tubes, and finally form hyphae.^9^ The small size and ubiquitous nature of these fungal spores requires that the human innate immune system must continuously defend against inhaled conidia. Epithelial cells are the likely source of first contact in the mammalian lung, but recruitment of alveolar macrophages and neutrophils quickly follows.^10^ Recognition by macrophages typically results in phagocytosis of conidia, while recruitment of neutrophils provides an additional level of antifungal defense.^11^ In immunocompromised patients, specifically those with neutropenia, these host defenses are impaired and result in unregulated fungal growth with dangerous clinical manifestations.^12^ Far less is known about the adaptive immune response to *A. fumigatus*, but CD4+ and CD8+ T cells have been shown to play important roles in antifungal defense.^13, 14^ The balance of host adaptive immune cell subsets appears to play an important role in susceptibility to fungal disease, as was recently shown for the accumulation of Th17 cells in ABPA, chronic obstructive pulmonary disease, and CF patients with acute airway inflammation.^15, 16^ Interestingly, tolerance to inhaled conidia was found to be mainly mediated by antigen-specific FOXP3+ regulatory T cells.^17^

In addition to *Aspergilli*, species of *Alternaria, Cladosporium*, and *Penicillium* are among the most commonly inhaled ascomycete molds capable of causing allergic diseases in humans.^2, 18^ Organisms in the genus *Penicillium* are genetically closely related to the *Aspergilli* and exhibit similar lifecycles; however, *Penicillium* species rarely cause infectious disease, but rather asthma and allergic rhinitis in adults and children.^19^ *Alternaria* and *Cladosporium* species are a major source of disease in plants, but they also represent important environmental allergens associated with asthma and asthma exacerbations.^3^ Due to their abundance in nature, it is not surprising that conidia and/or hyphal fragments containing immune-activating allergenic proteins from these fungi are thought to be a major source of fungal-induced allergy.^20^ Many allergens have been identified from each organism, with *A. fumigatus* containing 25 known allergen proteins, and *A. alternata, C. herbarum*, and *P. chrysogenum* (*rubens*) ranging from 6-17 protein antigens.^2, 21^ Despite this knowledge of allergens, our understanding of the complex interplay between fungus and host remains limited. The protein, lipid, and carbohydrate composition of a conidium has a direct impact on its recognition by the immune system and the following response of either tolerance or inflammation.^8, 13, 22^

The conidium of *A. fumigatus* is the best studied mold spore to date.^23^ Previous studies have elucidated the cell wall and surface proteome of *A. fumigatus* conidia to understand the molecules that likely interact with the host.^24–26^ Throughout germination the surface of the fungi changes dramatically, in terms of both the cell wall proteome and carbohydrate organization.^27, 28^ One major protein on the surface of conidia is the RodA hydrophobin, which encases resting conidia in a hydrophobic coat to minimize interactions with potential hazards. This hydrophobic rodlet layer is composed of a regular arrangement of the RodA protein interwoven amongst a layer of dihydroxynaphthalene (DHN)-melanin pigment. Upon encountering favorable growth conditions, resting conidia begin germination; a process that is highlighted by disassembly of the rodlet layer and degradation of the DHN-melanin polymer.^29–31^ Despite this key function, RodA is not required for virulence in a mammalian infection model.^30, 32^ Aside from RodA, the functions of few other *A. fumigatus* surface-exposed proteins are known. In many cases, these proteins seem to play important, but enigmatic roles in fungal biology, as is the case with the recently described CcpA protein, which serves as a conidial protein important for virulence in a mouse infection model.^24^ CcpA appears to have a structural role that normally masks other antigenic structures in the conidial cell wall. Despite our increasing understanding of the biology of fungal conidia, our ability to rapidly detect, diagnose, and treat fungal pathogens remains limited.^33^ Our capacity to detect airborne fungi remains particularly limited and slow, despite the obvious importance of fungal conidia in initiating invasive lung infections or allergic reactions in susceptible populations.^34–36^ As the burden of fungal pathogens in the clinic increases, improvements in each of these realms are desperately needed.^37^

In this study we further elucidated the surface-exposed proteome and secreted/shed proteins of allergy-inducing fungal conidia to reveal new fungal surface antigens as potential diagnostic or therapeutic targets. In addition, similarities and differences between the mold species were discovered, as well as the influence of environmental factors like temperature on the composition of the conidial surface proteome. We also focused on the biology of two particularly abundant, previously uncharacterized cell wall proteins of *A. fumigatus*, Afu4g09600 and Afu4g09310, which are denoted here as “<u>C</u>ell <u>W</u>all <u>E</u>nriched <u>A</u>” (CweA) and “<u>S</u>mall <u>C</u>ell <u>W</u>all Peptide <u>A</u>” (ScwA), respectively.

## Experimental Procedures

### Fungal strains and cultivation

All strains used in this study are listed in **Table S1**. Cultivation of *A. alternata* (Fries) Keissler (ATCC 66981), *P. rubens* WIS 54-1255 (ATCC 28089; formerly *P. chrysogenum WIS 54-1255*), and *A. fumigatus* CEA10 was performed on *Aspergillus* minimal medium (AMM; 70 mM NaNO_3_, 11.2 mM KH_2_PO_4_, 7 mM KCl, 2 mM MgSO_4_, and 1 μL/mL trace element solution (pH 6.5)) agar plates with 1% (w/v) glucose for 7 days at room temperature for the comparative analysis. The trace element solution was composed of 1 g FeSO_4_ • 7 H_2_O, 8.8 g ZnSO_4_ • 7 H_2_O, 0.4 g CuSO_4_ • 5 H_2_O, 0.15 g MnSO_4_ • H_2_O, 0.1 g NaB_4_O_7_ • 10 H_2_O, 0.05 g (NH_4_)_6_Mo_7_O_24_ • 4 H_2_O, and ultra-filtrated water to 1000 mL.^38, 39^ *C. herbarum* Link: Fries (ATCC MYA-4682) was cultivated on malt agar plates for 7 days at room temperature for the comparative analysis as AMM was not a permissible growth condition. All conidia were harvested in sterile, ultra-filtrated water without surfactants to minimize disruption of the surface proteome. Resting conidia from three 10-cm plates (~1×10^9^ conidia) were used immediately after collection for all trypsin shaving experiments. For the *A. fumigatus* germination experiment, resting conidia of *A. fumigatus* collected from three 10-cm plates grown for 7 days at 37°C were germinated with shaking in RPMI 1640 (Lonza) liquid media supplemented with glucose to a final concentration of 2% (w/v) at 37°C to produce swollen conidia (5 h), germlings (8 h), and hyphae (14 h). For the comparative analysis between *A. fumigatus, P. rubens, A. alternata*, and *C. herbarum*, resting conidia were prepared at room temperature for 7 days as described above and then transferred to 50 mL of RPMI media shaking (200 rpm) at 37°C for 5 hours in Erlenmeyer flasks. Fungal conidia were enumerated using a CASY^®^ Cell Counter (Omni Life Science) with care taken to resuspend thoroughly to avoid clumps.

### Extraction of surface proteins by trypsin shaving

Surface proteins were extracted as described previously.^24^ Briefly, resting conidia, swollen conidia, germlings, and/or hyphae of *A. fumigatus, A. alternata, C. herbarum*, and *P. rubens* were each washed twice with 25 mM ammonium bicarbonate and collected by centrifugation (1,800 × *g* for 10 min at room temperature). Samples were resuspended in 800 μL of 25 mM ammonium bicarbonate and treated with 5 μg MS-approved trypsin (Serva) for 5 min at 37°C with gentle agitation. Samples were then immediately passed through a 0.2 μm cellulose acetate filter (Sartorius) and collected in a microcentrifuge tube, followed by washing of the syringe filter with 200 μL of 25 mM ammonium bicarbonate. 9 μL of 89% (v/v) formic acid was added to stop the tryptic digestion. The samples were dried using a SpeedVac Concentrator (Thermo-Fisher), resuspended in 25 μL of 2% (v/v) ACN and 0.05% (v/v) TFA, and centrifuged for 15 min through a 10 kDa cutoff microfuge column (VWR Scientific). For LC-MS/MS method #2 described in detail below, the 10 kDa cutoff centrifugation step was replaced with centrifugation through a 0.22-μm-pore-size Spin X cellulose acetate spin filter (Corning Costar). Resting conidia and germinating conidia after 8 hours were collected after trypsin treatment and tested for lysis using a brief (15 min) staining with 40 μM propidium iodide followed by imaging with a Zeiss Axio Imager.M2 microscope (Zeiss).

### Purification of secreted proteins from swollen conidia

For the identification of secreted proteins, ~2×10^9^ resting conidia were germinated as described above for 5 h, before collection of the supernatant by filtration through a 0.22 μm filter and addition of TFA to a final concentration of 0.1% (v/v). Protein was purified by SPE with a Chromabond SPE-C4 cartridge (Macherey-Nagel, Germany). Briefly, columns were washed with 4 mL methanol and reconstituted with 4 mL 0.1% (v/v) TFA. Sample supernatants were aspirated through columns, washed with 2 mL 5% (v/v) MeOH / 0.1% (v/v) TFA, and eluted in 4 mL 0.1% (v/v) TFA 80/10 ACN/H_2_O (v/v). Samples were then dried by SpeedVac, resolubilized in 100 μL denaturation buffer (50 mM triethylammonium bicarbonate (TEAB) in 50/50 trifluoroethanol (TFE)/H_2_O (v/v)), sonicated for 15 min, and homogenized by Vortex mixing. Following 10 min at 90°C, samples were collected by centrifugation (1,800 × *g* for 10 min at room temperature) and cooled on ice. Protein content was measured with a Merck Millipore Direct Detect spectrometer. The samples were then reduced by adding 2 μL of reduction buffer (500 mM tris(2-carboxyethyl)phosphine in 100 mM TEAB) to 100 μL of sample (<1 μg/μL) for 1 h at 55°C. 2 μL of fresh alkylation buffer (625 mM iodoacetamide) was added to each sample and incubated for 30 min at room temperature, in the dark. Following evaporation by SpeedVac, the samples were resolubilized in 95 μL of 100 mM TEAB and sonicated for 10 min. The samples were treated with 2 μg of MS-approved trypsin (Serva) for 18 h at 37°C. The reaction was stopped with 10 μL of 10% (v/v) formic acid and again evaporated using the SpeedVac. Peptides were resolubilized in 25 μL 0.05% (v/v) TFA in 2/98 ACN/H_2_O (v/v) and sonicated for 15 min in a water bath before transfer to a 10 kDa molecular weight cut-off filter (VWR Scientific). After 15 min of centrifugation at 14,000 × *g* (20°C), the samples were transferred to HPLC vials and stored at −20°C.

### LC-MS/MS analysis

Three similar methods (denoted M#1, M#2, and M#3) were used for LC-MS/MS analysis due to advances in our proteomics approaches over time and differences in sample processing, *e.g*. trypsin shaving *versus* secretome analysis. A wild-type control was always included in each new analysis for comparison. LC-MS/MS analysis of trypsin-shaved surface peptides (M#1, M#2) and tryptic peptides of secreted proteins (M#3) were performed on an Ultimate 3000 RSLC nano instrument coupled to either a QExactive Plus (M#1, M#3) or an HF mass spectrometer (M#2; Thermo Fisher Scientific) as described previously.^24^ Tryptic peptides were trapped for 4 min on an Acclaim Pep Map 100 column (2 cm × 75 μm, 3 μm) at a flow-rate of 5 μL/min. The peptides were then separated on an Acclaim Pep Map column (50 cm × 75 μm, 2 μm) using a binary gradient (A: 0.1% (v/v) formic acid in H2O; B: 0.1% (v/v) formic acid in 90:10 (v/v) ACN/H2O).: M#1: 0-4 min at 4% B, 10 min at 7% B, 40 min at 10% B, 60 min at 15% B, 80 min at 25% B, 90 min at 35% B, 110 min at 50% B, 115 min at 60% B, 120-125 min at 96% B, 125.1-150 min at 4 % B. M#2: 0-4 min at 4% B, 5 min at 8% B, 20 min at 12% B, 30 min at 18% B, 40 min at 25% B, 50 min at 35% B, 57 min at 50% B, 62-65 min at 96% B, 65.1-90 min at 4% B. M#3: 0-5 min at 4% B, 30 min at 7% B, 60 min at 10% B, 100 min at 15% B, 140 min at 25% B, 180 min at 45% B, 200 min at 65% B, 210-215 min at 96% B, 215.1-240 min at 4% B. Positively charged ions were generated by a Nanospray Flex Ion Source (Thermo Fisher Scientific) using a stainless steel emitter with 2.2 kV spray voltage. Ions were measured in Full MS / dd MS2 (Top10) mode, *i.e*. precursors were scanned at m/z 300-1500 (R: 70,000 FWHM (M#1/#3)/: 60,000 FWHM (M#2); AGC target: 1·106e6; max. IT: 120 ms (M#1/#3) / 100 ms (M#2)). Fragment ions generated in the HCD cell at 30% normalized collision energy using N2 were scanned (R: 17,500 FWHM (M#1/#3) / 15,000 FWHM (M#2); AGC target: 2e5; max. IT: 120 ms (M#1/#3) / 100 ms (M#2)) in a data-dependent manner (dynamic exclusion: 30 s).

### Database search and data analysis of trypsin-shaved surface peptides

The MS/MS data were searched against the *A. fumigatus* Af293 genome of the *Aspergillus* Genome Database (AspGD); Uniprot Proteome ID UP000000724 for *P. rubens;* Uniprot Proteome ID UP000077248 for *A. alternata;* and the Joint Genome Institute MycoCosm annotation against the proteome of the related organism *C. fulvum* (described in^40, 41^) for *C. herbarum*, using Proteome Discoverer 1.4 and 2.2 and the algorithms of Mascot 2.4.1, Sequest HT, and MS Amanda as described previously.^24^ Briefly, we allowed for two missed cleavages for tryptic peptides, a precursor mass tolerance set to 10 ppm, and a fragment mass tolerance set to 0.02 Da. The dynamic modification was set as oxidation of Met. For secretome analysis (M#3) the static modification was set to carbamidomethylation of Cys. In all cases, at least 2 peptides per protein and a strict target false discovery (FDR) rate of < 1% were required for positive protein hits. In all cases, experiments were performed as three biological replicates with multiple technical replicates in the LC-MS/MS device. Proteins had to be identified in at least two of three replicates of a given condition to be included in the presented analyses, including supplemental tables.

## Data availability

The mass spectrometry proteomics data have been deposited to the ProteomeXchange Consortium via the PRIDE partner repository^42^ with the dataset identifier PXD017067.

### Genetic manipulation of *A. fumigatus*

All oligonucleotides utilized in this study are listed in **Table S2.** The genetic deletions of *cweA* (Afu4g09600) and *scwA* (Afu4g09310) were generated using a 3-fragment PCR method, as previously described.^43^ ScwA, a small protein, was reannotated in *A. fumigatus* CEA10 as indicated in **Figure S1A** (V. Valiante, personal communication). For the knockout constructs, flanking regions of ~1 kb were amplified for each gene using *A. fumigatus* CEA17 genomic DNA as template and the appropriate primer pairs (p1/p2 and p3/p4 indicated in **Figure S1B, C**). The pyrithiamine resistance marker (*ptrA*) was amplified from the pSK275 plasmid using primers ptrA-F/R. Using Phusion polymerase (Thermo Scientific) and equimolar amounts of the three fragments as template, primers p1 and p4 directed amplification of the entire construct (**Figure S1D**). Knockouts of *A. fumigatus* were generated by homologous recombination following the transformation of protoplasts.^44^ Knockouts were verified by PCR and Southern blot according to standard procedures as described below (**Figure S1E, F**).

Complemented strains were produced by knocking in the gene of interest to the endogenous locus of the corresponding knockout strain (**Figure S1G-J**). Constructs were produced by fusion PCR using primers listed in **Table S2**. Briefly, to complement strains CEA17Δ*scwA and* CEA17Δ*cweA*, plasmids *pUC19_scwAc* and *pUC19_cweAc* were produced and transformed into protoplasts as previously described.^44^ Both plasmids were generated using a Gibson cloning and assembly kit (BioLabs) according to the manufacturer’s instructions. For *pUC19_scwAc*, the DNA fragments were generated as follows: the *scwA* gene and its 3’ flanking region were PCR amplified using primers scwA_F/scwA_hph_R and scwA_hph_F/scwA_R, respectively. The hygromycin resistance cassette (*hph*) was amplified from the *pJWx41* plasmid with primers hph_scwA_F/hph_scwA_R, and the plasmid *pUC19* backbone was amplified by PCR using primers pUC19_F/pUC19_R. The construct from transformation was amplified using primer pair scwA_F and scwA_R. For *pUC19_cweAc*, the *cweA* gene and its 3’ flanking region were PCR amplified using primers cweA_F/cweA_hph_R and cweA_hph_F/cweA_R, respectively. The hygromycin resistance cassette (*hph*) was amplified from *pJWx41* plasmid with primers hph_cweA_F/hph_cweA_R, and the plasmid *pUC19* backbone was amplified by PCR using primers pUC19_F/pUC19_R. Primer pair cweA_F/cweA_R was used for amplification of the construct for transformation.

### Southern blot analysis

Southern blots to validate genetic deletion were carried out according to standard protocols.^45^ All sequence information was obtained from the Ensembl Fungi Database.^46^ Genomic DNA of *A. fumigatus* was isolated using the MasterPure Yeast DNA Purification Kit (Epicentre). Genomic DNA of *A. fumigatus* was digested with the appropriate restriction enzymes (New England Biolabs). DNA fragments were separated on an agarose gel followed by blotting onto Hybond N+ nylon membranes (GE Healthcare Bio-Sciences). DNA probes were amplified with digoxigenin-UTP (DIG-UTP) in the PCR reaction using primer pairs cweA-1/2, scwA-3/4, cweA_hph_F/cweA_R, and scwA_F/scwA_hph_R for Δ*cweA*, Δ*scwA*, *cweAc*, and *scwA*c, respectively (primers in Table S2). Genomic DNA of candidates Δ*cweA*, Δ*scwA*, *cweAc*, and *scwA*c was digested using restriction enzymes *Cla*I, *Hind*III, *Bam*HI, and *Fco*RI, respectively, hybridized to the appropriate probe in DIG Easy Hyb (Roche Applied Science), and detected using CDP-Star ready-to-use kit (Roche Applied Science), according to the manufacturers’ recommendations. Chemiluminescence signals were detected by a Fusion-FX CCD camera (Vilbert Lourmat).

### Growth Assays of *A. fumigatus* knockouts

Wild-type and knockout conidia were collected in water from AMM agar plates after 5 days of growth. Conidia were counted using a CASY^®^ Cell Counter (Omni Life Science), diluted in droplet solution (0.9% (v/v) NaCl, 0.1% (v/v) Agar, 0.01% (v/v) Tween 20) and added to plates in 5 μL drops containing 10^2^, 10^3^, 10^4^, or 10^5^ conidia. Growth was monitored over time at 37°C and images were collected before overgrowth of the plate. Cell stress was induced using common stressors^47, 48^ including caffeine (Fluka), caspofungin acetate (Cancidas), calcofluor white (Sigma), congo red (Sigma), and SDS (Serva) all dissolved in water (except for calcofluor white, which was dissolved in Tris-HCl) and added directly to cooling agar at the concentrations denoted in the text. Plates were prepared fresh for each experiment and growth was allowed to continue for 72 hours. Germination assays were performed as above by microscopically determining the ratio of spores undergoing germination (germ-tube formation) at each time point.

### Bioinformatic analyses

Phylogenetic analyses were preformed using sequences gathered from FungiDB^49^ for each ortholog of *A. fumigatus* CweA and ScwA. Additional protein-protein BLAST (blastp) searches were performed to confirm the collection of all orthologs. Alignments and phylogenetic trees were then produced using the DNAStar MegAlign software package (DNASTAR Inc., USA) using a ClustalW alignment.

The InParanoid algorithm was used to identify ortholog clusters for pairwise species comparison.^50^ The default algorithm parameters were used: a score cut-off of 40 bits, an overlap cut-off of 50%, and a segment cut-off of 25%. The proteins with an ortholog score <0.95 contained in the ortholog clusters were excluded from the analysis. “One to many” and “many to many” orthologous relationships were included in the ortholog clusters. An ortholog group among 3 or 4 species was found only if the ortholog group was identified for all the pairings. Furthermore, the InParanoid-defined ortholog clusters with at least one expressed protein were also identified.

## Results and Discussion

### The *A. fumigatus* surface proteome is dynamic across germination

In this study, we sought to expand the repertoire of *A. fumigatus* conidial surface and secreted proteins that potentially trigger allergy or promote infection and compare the proteins exposed on the surface of *A. fumigatus* conidia with other important allergenic molds. Initially, we utilized a previously-described trypsin-shaving proteomics approach^24^ to identify surface-exposed proteins throughout germination of the filamentous fungus, *A. fumigatus* strain CEA10 (**Figure 1A**). Samples were collected from resting (0 h) conidia grown at 37°C on AMM agar plates, and swollen conidia (5 h), germlings (8 h), and hyphae (14 h) germinated at 37°C in RPMI liquid media. We identified 177 unique proteins across all germination time points. 75/177 proteins had a predicted signal peptide cleavage site using the SignalP program,^51^ 15/177 contained at least one predicted transmembrane helix using TMHMM Server v. 2.0 (services.healthtech.dtu.dk),^52^ and 13/177 contained putative GPI anchors using PredGPI (**Table S3**;^53^). When compared to previous trypsin-shaving data from the literature, 43/55 proteins from CEA10 were found on the surface of *A. fumigatus* strain D141 resting conidia under similar growth conditions, whereas in swollen conidia, 98/139 CEA10 proteins were on the surface of D141 swollen conidia (**Figure 1B-C**;^24^). These results suggest that the surface proteome of different *A. fumigatus* strains exhibit high similarities under standard growth conditions. The differences still observed could be due to slight differences in growth media. The surface proteins identified have a diverse array of cellular localizations and activities using Gene Ontology Enrichment (fungidb.org). These include localization to the cell wall (72/177; GO:0005618), cytoplasm (71/177; GO: 0005737), and external encapsulating structure (72/177; GO:0030312) with activities related to catalytic activity (89/177; GO:0003824), binding (78/177; GO:0005488), oxidoreductase activity (22/177; GO:0016491), and nucleic acid binding activity (16/177; GO:0003676), among others.

**Figure 1.**
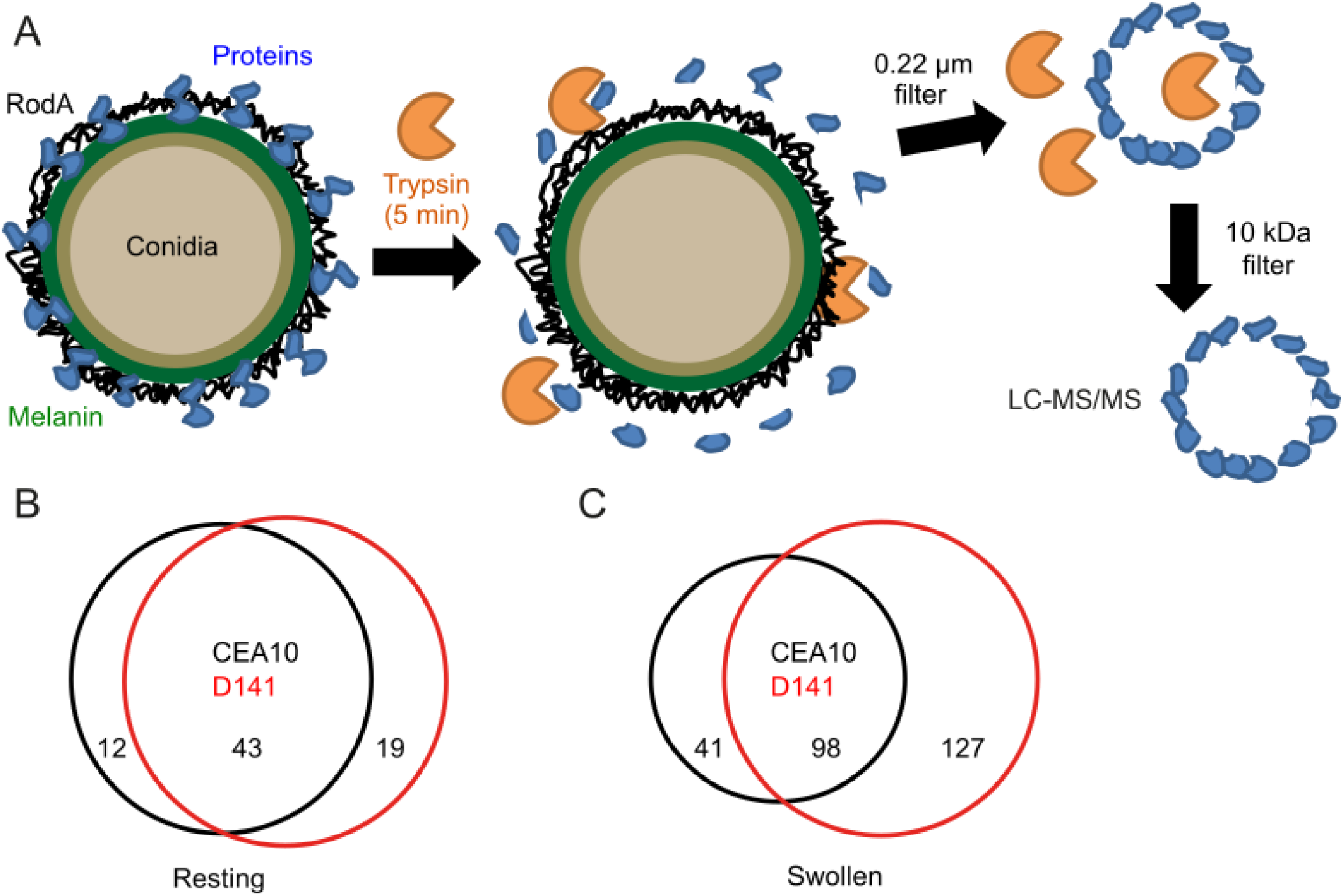
Trypsin-shaving proteomics of allergy-inducing fungi. A) Model of trypsin-shaving proteomics approach to identifying surface-exposed peptides. Scaled Venn diagrams produced using the Biovenn software^54^ show intersection of proteins identified by trypsin shaving of B) resting and C) swollen conidia from *A. fumigatus* strain CEA10 with D141 (D141 data described in^24^). Trypsin-shaving proteomics was performed as three separate biological replicates with multiple technical replicates measured by LC-MS/MS. To be included in the analysis, proteins had to be identified in at least two of three biological replicates.

Over the course of germination, 55 proteins were identified on resting conidia (see **Table 1** for Top 20), 140 on swollen conidia, 56 on germlings, and 109 on hyphae. In direct comparisons between samples, 35 proteins were found in all four stages of the germination time course, including the conidial hydrophobin, RodA (**Figure 2**). RodA exhibited the highest levels of normalized abundance throughout germination; ranging from 16.8% to 7.7% of the normalized peptide spectrum matches (PSMs) per length and consistent with previous study.^26, 32^ The majority of conidia underwent germination in our assays; however, a small fraction of conidia failed to germinate, which likely contributed to the abundance of RodA at later time points. In agreement, localization studies with a polyclonal RodA antibody revealed the presence of RodA on the surface of conidia, in biofilm cell walls, and in phialides during sporulation.^32^

**Figure 2.**
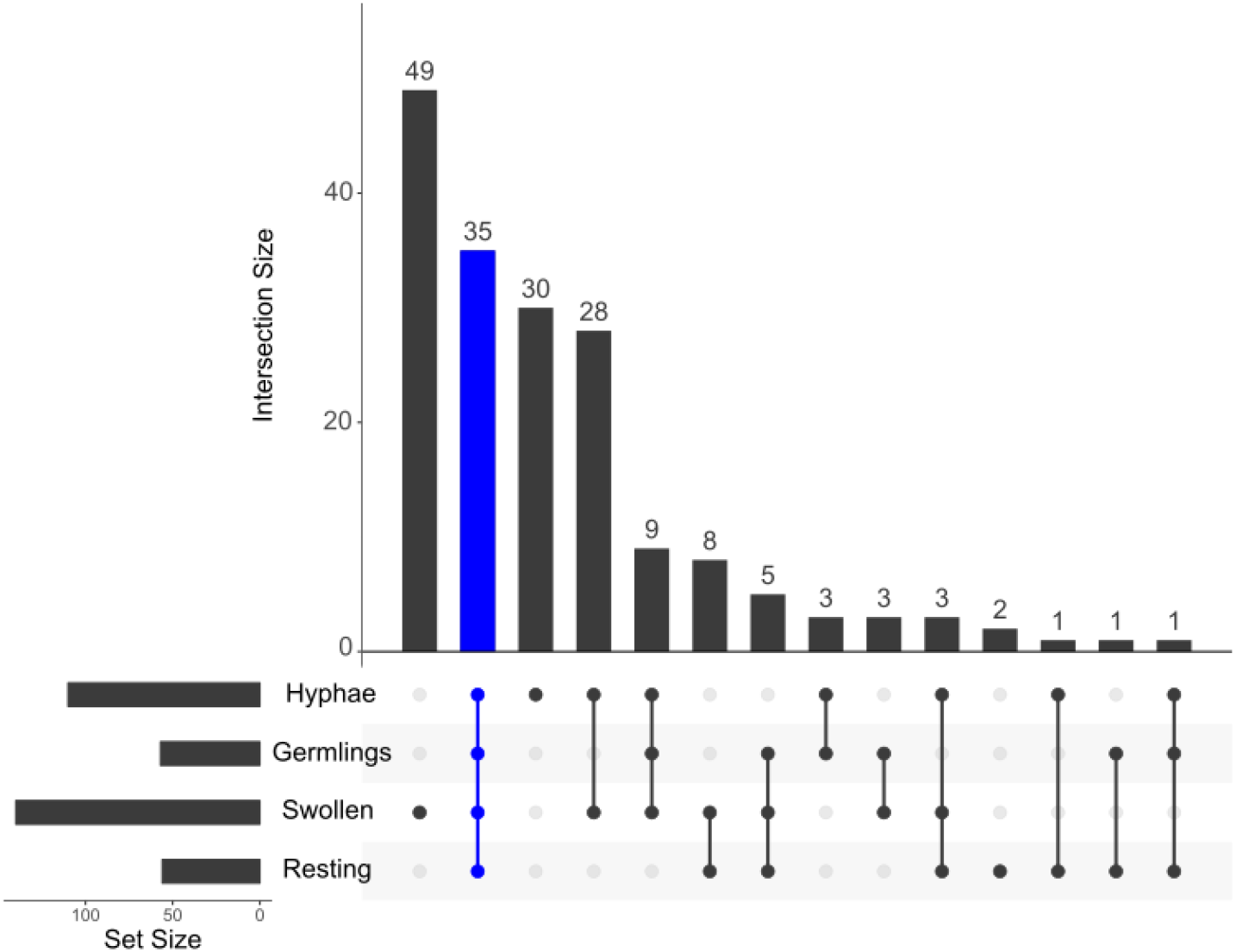
Trypsin shaving proteomics elucidates surface proteome of *A. fumigatus* conidia after germination in RPMI. 4-way comparison indicating the intersection of surface-exposed proteins from resting (0 h) and swollen conidia (5 h), germlings (8 h), and hyphae (14 h). The experiment was performed as three separate biological replicates with multiple technical replicates measured by LC-MS/MS and to be included in the analysis, proteins had to be identified in at least two of three biological replicates. Plot produced using the R package UpSet.

**Table 1.** Top 20 *A. fumigatus* proteins identified on resting conidia ranked by normalized spectral abundance factor (NSAF).

| Accession Number | Protein Name | NSAF | Description |
| --- | --- | --- | --- |
| Afu5g09580 | RodA | 0.126 | Conidial hydrophobin |
| Afu6g03210 | ConJ | 0.093 | Uncharacterized protein |
| Afu2g09030 | DppV | 0.055 | Secreted dipeptidyl-peptidase |
| Afu6g14470 | Afu6g14470 | 0.045 | Uncharacterized protein |
| Afu2g05635 | Afu2g05635 | 0.041 | Uncharacterized protein |
| Afu1g13670 | CcpA | 0.040 | Conidial cell wall protein |
| Afu4g09310 | Afu4g09310 | 0.037 | Uncharacterized protein |
| Afu4g09280 | Afu4g09280 | 0.033 | Uncharacterized protein |
| Afu5g14210 | Grg1 | 0.033 | Glucose-repressible gene |
| Afu1g14560 | MsdS | 0.030 | Putative 1,2-alpha-mannosidase |
| Afu8g01980 | Afu8g01980 | 0.027 | Uncharacterized protein |
| Afu5g03540 | Afu5g03540 | 0.024 | Putative flavin-linked sulfhydryl oxidase activity |
| Afu2g17530 | Abr2 | 0.023 | Laccase |
| Afu8g05020 | NagA | 0.022 | Putative secreted N-acetylhexosaminidase |
| Afu3g07160 | Afu3g07160 | 0.020 | Putative family 18 chitinase |
| Afu2g00680 | Afu2g00680 | 0.020 | Uncharacterized protein |
| Afu1g13550 | Afu1g13550 | 0.019 | Uncharacterized protein |
| Afu4g09600 | Afu4g09600 | 0.018 | Putative glycosphosphatidylinositol (GPI)-anchored cell wall protein |
| Afu4g01290 | Csn | 0.016 | Glycosyl hydrolase family 75 chitosanase |
| Afu5g09240 | Sod1 | 0.016 | Cu/Zn superoxide dismutase |

In addition to RodA, the other most prevalent proteins on the surface of resting conidia, as determined using normalized spectral abundance factor (NSAF;^55, 56^), were the protein of unknown function, ConJ, Afu6g14470, and Afu2g05635; the serine protease, DppV; and the previously described surface protein CcpA. ConJ shows high similarity to the conidiation-specific, light-induced protein 10 of *Neurospora crassa* and its transcripts are upregulated during early conidial development and murine infection with *A. fumigatus*.^26^ Afu6g14470 and Afu2g05635 remain relatively unstudied, but each contains a signal peptide cleavage site according to SignalP,^51^ and Afu6g14470 was previously detected by 2D-gel electrophoresis on the conidial surface of *A. fumigatus*.^25^ The protein DppV is a secreted dipeptidyl-peptidase and nonclassical serine protease^57^, which is known to be targeted by the T cell response in mice surviving *A. fumigatus* infection.^58^ The surface of swollen conidia exposed more proteins (140 unique proteins identified) than what was found on the surface of the other time points, which is consistent with previous studies comparing resting and swollen conidia.^24^ It is possible that reorganization of the conidial surface during germination results in the temporary availability of cell wall proteins to trypsin cleavage in some circumstances. The surface proteome of germlings was less complex, with no proteins unique to only germlings, suggesting that perhaps there is a restriction to the surface proteome in germlings. In hyphae, we again see slightly higher numbers of surface-exposed proteins, with 30 proteins found that were not identified in the other samples (**Figure 2**).

### Knockout of abundant conidial surface proteins does not alter surface proteome

We were interested in further studying several of the more highly abundant surface proteins from *A. fumigatus*. The putative GPI-anchored cell wall protein CweA (Afu4g09600, 980 amino acids) and the protein ScwA (Afu4g09310, 93 amino acids) were both predicted to encode a signal peptide for entry into the secretory pathway (SignalP), but otherwise lacked recognizable conserved domains or characterized orthologs. In addition to being highly abundant, CweA and ScwA were observed on the surface throughout germination by trypsin shaving (**Table S3**) and were previously found on the conidial surface in other studies.^24, 59^ Interestingly, *cweA* contains multiple coding tandem repeats, and its transcript was shown to be up-regulated upon membrane stress.^60^

Since ScwA is such a small protein, we first confirmed the annotation of the genetic loci in the CEA10 genome by multi-sequence alignment to related organisms after a resequencing of CEA10 *scwA* (**Figure S1A**). ScwA appears to exclusively belong to a small subset of the *Aspergillus* genus (**Figure S2A**), while CweA is widely found in both the *Aspergillus* and *Penicillium* genera (**Figure S2B**), with an ortholog expressed in *P. rubens*. Genetic deletions of *cweA* (Afu4g09600) and *scwA* (Afu4g09310) were produced in *A. fumigatus* CEA17 by homologous recombination following transformation of protoplasts (**Figure S1B-F).** Both knockouts grew normally on AMM and malt agar plates (**Figure S3A**). The genetic deletion of *cweA* provided a growth advantage relative to wild type on Congo Red agar plates, a known cell wall stressor,^47, 48^ whereas *ΔscwA* exhibited only a minor growth inhibition (**Figure 3A**). This finding is consistent with data from the literature showing that *cweA* was downregulated in response to Congo Red stress.^60^ Under calcofluor white stress, the *ΔscwA* strain exhibited a slight advantage, whereas *ΔcweA* exhibited a more robust growth advantage, all of which could be complemented (**Figure 3A**). Challenge with other cell wall and membrane stressors, including caffeine, caspofungin, and SDS revealed no obvious defects by droplet growth inhibition assay (**Figure S3B**). Taken together, these results suggest that the abundant surface proteins ScwA and CweA are important for fine tuning the conidial response to stress.

**Figure 3.**
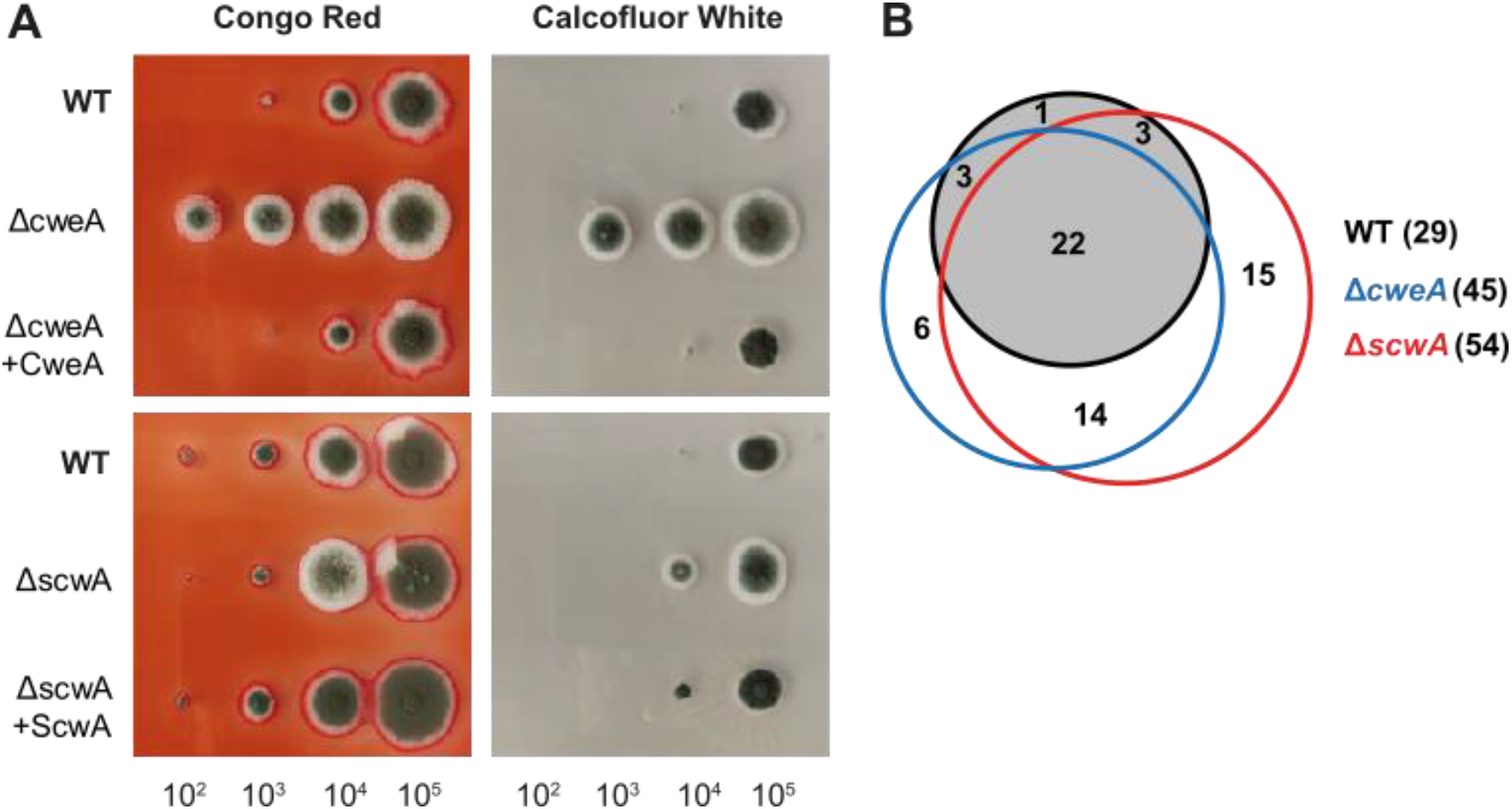
Phenotypic analysis of *A. fumigatus* conidial surface proteins. A) Droplet growth inhibition assays for wild-type, Δ*cweA*, and Δ*scwA* knockout strains grown on AMM agar plates supplemented with 35 μg/mL Congo red or 50 μg/mL calcofluor white. Images are representative of three separate experiments. B) Scaled Venn diagram indicating intersection of surface proteins identified by trypsin shaving of wild-type, Δ*cweA*, and Δ*scwA* knockout strains. Trypsin-shaving proteomics was performed as three separate biological replicates with multiple technical replicates measured by LC-MS/MS, and to be included in the analysis, proteins had to be identified in at least two of three biological replicates.

Our previous characterization of the surface protein CcpA revealed an altered conidial surface proteome after growth on AMM agar plates.^24^ In this case, it was postulated that CcpA functions to mask other cell wall proteins or other surface molecules that are normally hidden in the conidial cell wall. To explore if other cell wall protein knockouts resulted in a similar alteration to the surface proteome of resting conidia, we performed trypsin-shaving on the Δ*cweA* and Δ*scwA* knockout strains compared to the wild-type CEA17 *akuB^KU80^* strain used to create the knockouts.^61^ In both cases, we saw very little change to the exposed surface proteome, suggesting that the function of CcpA in masking surface-exposed proteins is not a common feature of all abundant surface proteins, but rather a specific feature of CcpA (**Table S4; Figure 3B**). Ultimately, these results suggest that the cell surface is quite robust and adaptable, as even the removal of abundant cell surface proteins like CweA and ScwA yield few major phenotypes and little changes in the overall structure. Further study will be required to fully understand the redundancy of conidial surface proteins in cell wall maintenance and stress response.

### Growth time and temperature dictate the surface proteome of *A. fumigatus* conidia

As described above, the surface proteome undergoes dynamic changes during the germination process. Previous studies showed that environmental conditions during conidiation affect the stress tolerance of *A. fumigatus* conidia.^62^ We thus wanted to understand if and how the temperature during conidiation influences the surface proteome of *A. fumigatus* resting conidia. We performed trypsin-shaving proteomics on resting conidia grown for 7 days at room temperature, 37°C, or 42°C and for 14 days at room temperature or 37°C. Surprisingly, conidia produced at room temperature and 42°C had more surface-exposed proteins than spores grown at 37°C after 7 days of growth (**Figure 4A, Table S5**). After 14 days of incubation, conidia grown at room temperature and at 37°C both exhibited far fewer proteins on the surface, 19 and 29, respectively (**Figure 4B, Table S5**). These results suggest that the surface proteome of conidia is potentially restricted or degraded over developmental time. In agreement with this idea of a surface proteome restriction is previous work showing that the whole conidial proteome has a lower complexity at 15 days compared to conidia after 3 or 30 days of growth.^63^ It remains unclear how changes to the intracellular proteome might influence the surface proteome during conidial development, but one explanation is that more cytosolic proteins contaminate the conidial surface at 7 days compared to 14 days, at which time additional proteolytic degradation has occurred. It is also possible that as the RodA coat and melanin layers become more contiguous during development, it is harder for trypsin to penetrate into the cell wall to release surface-exposed peptides. Similarly, increases in hydrophobicity during development may result in decreased ability of the trypsin protease to reach the surface. We hope that future studies will be able to better define the exact mechanism by which the surface proteome is limited during development.

**Figure 4.**
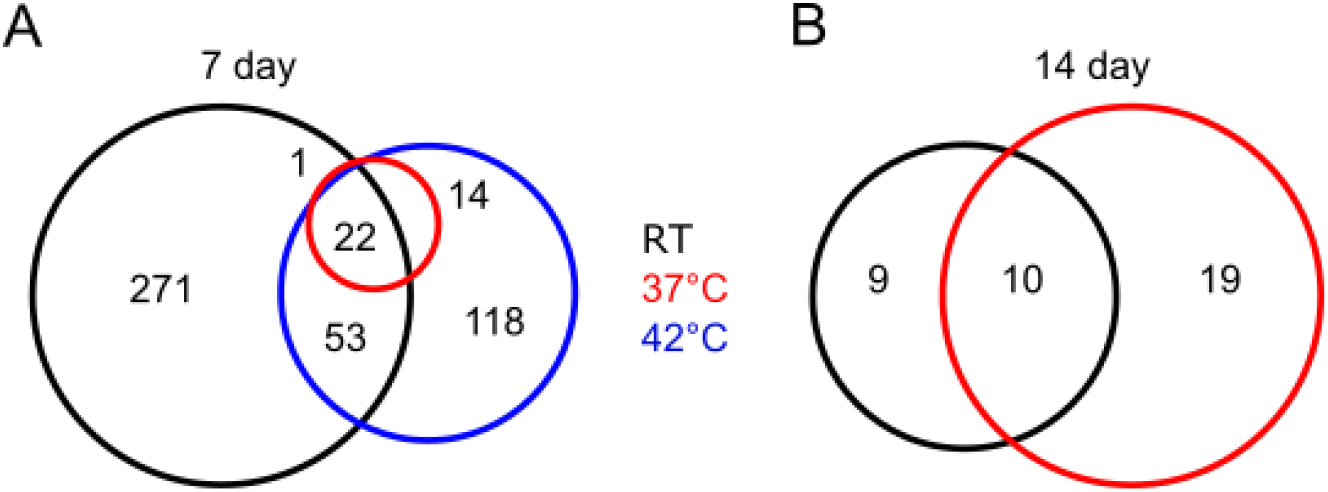
Temperature and time influence the *A. fumigatus* surface proteome. A) Scaled Venn diagrams show the intersection of proteins identified by trypsin shaving on the surface of resting conidia after A) 7 days at 37°C, 42°C, or room temperature or B) 14 days at 37°C or room temperature. Experiments were performed in triplicate with multiple technical replicates and to be included in the analysis, proteins had to be identified in at least two of three biological replicates.

Interestingly, the ScwA protein appears to be only on the surface of conidia grown at 37°C, and not room temperature, suggesting a possible role in temperature adaptation. However, we see no obvious growth difference for the knockout strain at 37°C or room temperature. Five proteins were identified on resting conidia grown at all tested time points and temperatures (**Table 2**), including one previously uncharacterized protein, Afu8g01980, which is a putative phosphotriesterase containing a TolB-like six-bladed beta-propeller domain (https://www.ebi.ac.uk/interpro/).^64^ Overall, these results have obvious implications for the study of *A. fumigatus* pathogenesis and reaffirm that the growth temperature of conidia can have a drastic effect on surface protein expression.^62^ One partial explanation for the observed temperature-dependent effects is the difference in growth rate of *A. fumigatus* on agar plates at room temperature and 37°C, where sporulation requires several additional days at room temperature. At 14 days of incubation the conidia are mature under both conditions, and at this time point we observe a similar number of proteins on the surface of conidia (**Figure 4B**). Despite a comparable number of surface proteins, 19 for spores grown at room temperature and 29 for spores cultivated at 37°C, the composition of the surface proteome is not the same at 14 days (10 shared proteins), suggesting some temperature-dependent regulation of surface-exposed proteins.

**Table 2.** *A. fumigatus* proteins identified on resting conidia at all tested time points and temperatures.

| Accession Number | Protein Name | Description |
| --- | --- | --- |
| Afu5g09580 | RodA | Conidial hydrophobin |
| Afu1g07440 | Hsp70 | Molecular chaperone |
| Afu8g01980 | Afu8g01980 | Uncharacterized protein |
| Afu2g17530 | Abr2 | Laccase |
| Afu1g05630 | Afu1g05630 | 40S ribosomal protein S3, putative |

### Comparative proteomics uncovers proteins on the surface/shed from allergenic fungal conidia

To refine our understanding of the surface-exposed cell wall proteins that might contribute to the initiation of allergic inflammation in the lung, we leveraged the power of comparative proteomics. To this end, the surface-exposed proteomes of 7-day-old resting conidia of *A. fumigatus* (CEA10) and three other prevalent allergy-inducing fungi, *A. alternata* (ATCC 66981), *C. herbarum* (ATCC MYA-4682), and *P. rubens* (ATCC 28089) grown at room temperature were assessed by trypsin-shaving proteomics. We first confirmed that the short trypsin treatment did not result in cell lysis in all cases for resting conidia and germlings by staining with the cell-impermeable dye propidium iodide (**Figure S4**). *A. alternata* and *C. herbarum* conidial populations contained a noticeable fraction of dead cells, which were permeable to propidium iodide in both the presence and absence of trypsin (**Figure S4**). Therefore, we believe that these datasets provide more information about trypsin-accessible proteins (and likely immune-accessible proteins), and less information about the exact surface composition of *A. alternata* and *C. herbarum*. Unfortunately, an annotated genome of *C. herbarum* is not available, so we relied on the mapping of peptides to the annotated *C. fulvum* genome for these analyses, which potentially results in the omission of some *C. herbarum-specific* surface proteins. *C. herbarum* was also unable to sporulate on AMM agar plates, and thus some of the observed differences may be due to differences between growth media. After growth at room temperature, conidia were collected and transferred to RPMI media at 37°C. This temperature and growth substrate switch was designed to better mimic the likely change in environment from growth outdoors or in a hospital environment to that of the lung of an immunocompromised patient.

LC-MS/MS analysis revealed 252 surface-exposed proteins on resting conidia of *A. fumigatus*, 473 on *A. alternata*, 240 on *C. herbarum*, and 254 on *P. rubens* (**Figure 5A; Table S6-S9**). On swollen conidia, we observed 49 surface-exposed proteins on *A. fumigatus*, 1,007 on *A. alternata*, 121 on *C. herbarum*, and 672 on *P. rubens. A. fumigatus* conidia harvested from cultures grown at room temperature again exhibited more proteins on the surface than conidia obtained from cultures grown at 37°C (**Figure 5B**). These results are also consistent with previous studies that link fungal growth variability to cultivation temperature,^62^ and again highlight the importance of selecting appropriate and consistent growth conditions. Interestingly, we observed conidia from mycelia grown at room temperature or 37°C for 7 days responded differently to germination in RPMI, such that swollen conidia exhibited vastly different surface proteomes after 5 hours at 37°C (**Figure 5C**). In particular, swollen conidia grown initially at room temperature responded by restricting their surface proteomes, whereas conidia grown initially at 37°C expanded their surface proteomes. The mechanism of this phenotype remains unknown but provides an interesting view of the dynamics of the surface proteome.

**Figure 5.**
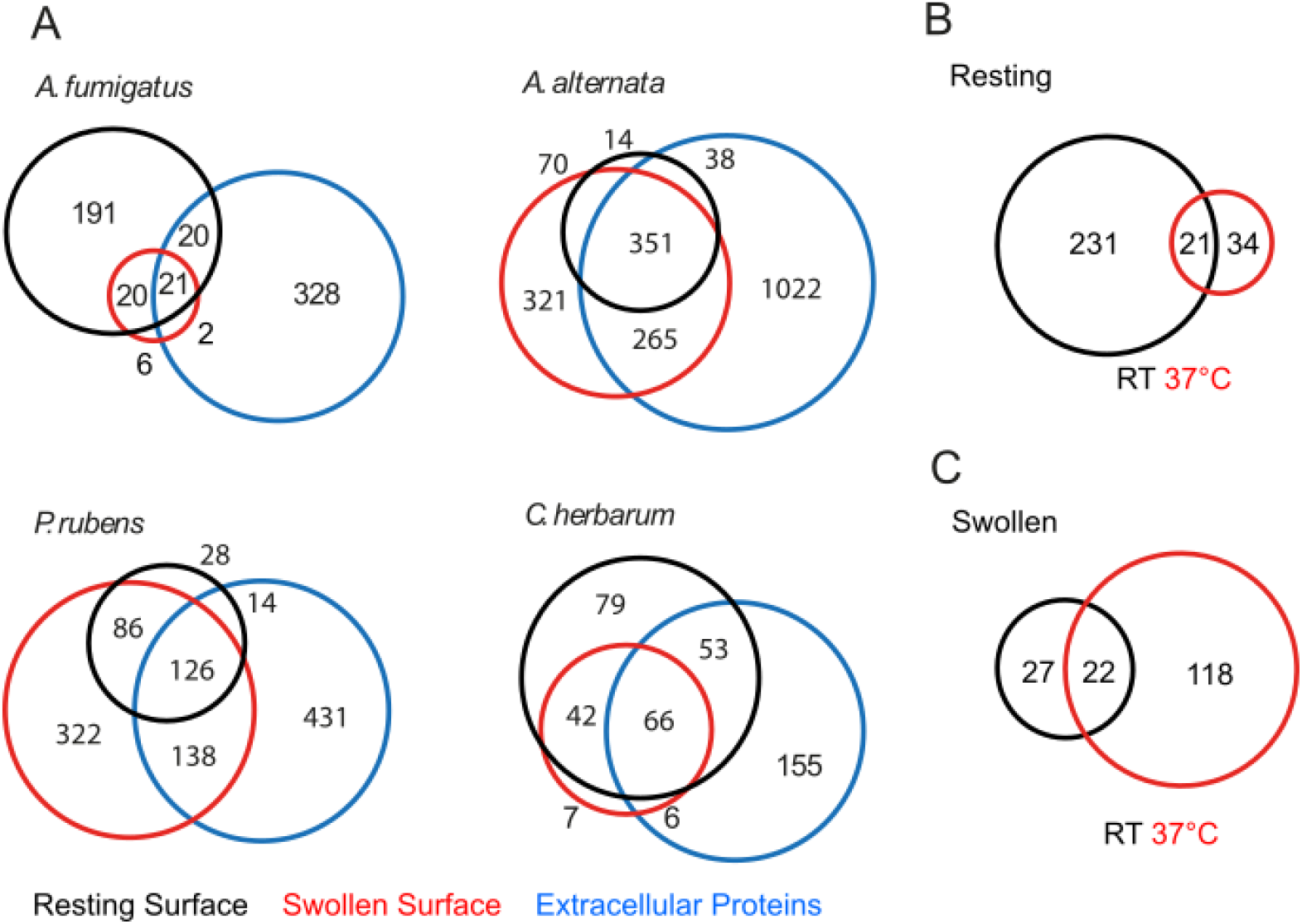
Comparative analysis reveals conserved surface proteins. A) Scaled Venn diagrams show the intersection of surface-exposed proteins of resting conidia and swollen conidia (5 h) and extracellular proteins (5 h) from *A. fumigatus, P. rubens, A. alternata*, and *C. herbarum* (annotated using *C. fulvum* genome) germinated at 37°C in RPMI. Although the experimental setup for *A. fumigatus* CEA10 grown at room temperature for 7 days shown in Figures 5A (black circle) was the same as that described in Figure 4A (black circle), it was redone for this experiment as an additional control. B) Scaled Venn diagrams indicate difference in surface proteome between *A. fumigatus* resting conidia isolated from mycelia grown at room temperature or 37°C. C) Scaled Venn diagrams indicate differences in the surface proteome between *A. fumigatus* swollen conidia isolated from mycelia grown at room temperature or 37°C. Experiments were performed in triplicate with multiple technical replicates and to be included in the analysis, proteins had to be identified in at least two of three biological replicates.

In addition to surface-exposed proteins, we were also interested in proteins secreted from swollen conidia that may serve as allergens. Our previous proteomics analyses revealed an abundance of *A. fumigatus* proteins available for trypsin cleavage on swollen conidia.^24^ To test the hypothesis that a subset of these newly surface-exposed proteins might also be secreted or shed from conidia, we performed solid-phase extraction of spent cell culture supernatant after 5 h of germination. Using this method, we were able to identify 371 *A. fumigatus* proteins secreted/shed after 5 h of incubation, 1,676 from *A. alternata*, 280 from *C. herbarum*, and 709 from *P. rubens* (**Figure 5A, Table S6-S9**). These findings highlight that many proteins exposed on the surface of germinating conidia are also secreted/shed into the local environment, where they may serve as potential allergens to activate or inhibit host responses. In fact, 13 known allergens were identified in *A. fumigatus*, including Asp f 1, Asp f 9, and Asp f 12. Of these 13 allergens, 9 were found only secreted/shed during swelling, two allergens were specific to resting conidia (Asp f 12, Asp f 23), and one was detected in all three conditions (Asp f 1). Three known allergens were identified in *A. alternata*, with two on the surface of resting and swollen conidia as well as secreted/shed from swelling conidia (Alt a 6, Alt a 7), and one found only secreted/shed from swelling conidia (Alt a 1). Four allergens were identified in analyses of *P. rubens:* Pc20g10360 (Beta-hexosaminidase), Pc21g16950 (transaldolase), and Pc21g16970 (vacuolar serine proteinase) were found on resting conidia, swollen conidia, and secreted/shed from swelling conidia; and the protein of unknown function Pc12g10650 was found only secreted/shed from swelling conidia.

Fungal secreted proteases are known to drive allergic airway diseases by hydrolyzing the glycoprotein complex fibrinogen (FGB), which leads to the activation of a TLR4-mediated immune response *via* the integrin Mac-1.^65^ The secreted serine protease, Alp1 (Asp f 13) of *A. fumgiatus* is a known promoter of airway hyperresponsiveness.^66^ Our proteomic data reveal that Alp1 is secreted from swollen conidia of *A. fumigatus*, like the serine protease Pc21g16970 of *P. rubens* (40% sequence similarity). No orthologs of Alp1 were found in the secretome of *A. alternata* or *C. herbarum*. However, *A. fumigatus* secretes an arsenal of other peptidases (Afu3g08290, Afu3g11400, Afu5g04330, AFu7g05930) whose orthologs are also detectable in the secretomes of all other investigated molds. All have the potential to initiate allergic and antifungal immunity.

### Ortholog analysis reveals a conserved surface proteome among allergenic fungi

Next, we used the InParanoid software package to assign ortholog groups between the different fungi in our comparative proteomics datasets.^50, 67^ The aim was to identify proteins that were conserved across all four organisms as well as to define proteins that were specific to one organism. We were able to identify 125 orthologous surface/secreted protein groups detected in all four organisms (**Table 3, Table S10**). In *A. fumigatus*, these orthologs corresponded mostly to proteins in the cytosol (91/125; GO:0005737), but a small subset was found in the extracellular region (37/125; GO:0005576) by Gene Ontology Enrichment (fungidb.org). The large number of proteins from the cytosol is indicative of the prevalent contamination of the cellular surface with proteins from the intracellular space by each of the four tested fungi. Many represent ribosomal proteins, chaperones, and metabolic enzymes, some of which have been shown to exhibit antigenic activity.^68^ It is possible that some of these proteins have moonlighting functions in the extracellular environment, as has been observed in other systems.^8^ For example, Asp f 22 (enolase) was recently shown to bind human plasma complement proteins to damage epithelial cell layers.^69^ Interestingly, 42 *A. fumigatus* proteins were detected in this experiment that had no ortholog in the related allergy-inducing species, including the Asp f 1 allergen. 102 proteins were specific to the extracellular space of *P. rubens*, 206 proteins unique to *A. alternata*, and 9 to *C. herbarum* (**Table S10**). Of note, we did observe the *P. rubens* orthologs of *A. fumigatus* CweA and CcpA on *P. rubens* conidia, perhaps suggesting a conserved function for these proteins in *P. rubens*.

**Table 3.** Ortholog analysis on surface/secreted proteins of allergy-inducing fungi grown at room temperature. (See also Table S10 in the supplements)

| Organism A | Organism B | Organism C | Organism D | Total Ortholog Groups from detected proteins | Ortholog Groups w/ orthologs detected in all organisms |
| --- | --- | --- | --- | --- | --- |
| *A. fumigatus* | *P. rubens* | *C. herbarum* | *A. alternata* | 1617 | 125 |
| *A. fumigatus* | *P. rubens* | *C. herbarum* | - | 1050 | 134 |
| *A. fumigatus* | *P. rubens* | - | *A. alternata* | 1773 | 289 |
| *A. fumigatus* | - | *C. herbarum* | *A. alternata* | 1589 | 146 |
| - | *P. rubens* | *C. herbarum* | *A. alternata* | 1657 | 258 |
| *A. fumigatus* | *P. rubens* | - | - | 1146 | 351 |
| *A. fumigatus* | - | *C. herbarum* | - | 664 | 156 |
| *A. fumigatus* | - | - | *A. alternata* | 1737 | 377 |
| - | *P. rubens* | - | *A. alternata* | 1835 | 726 |
| - | *P. rubens* | *C. herbarum* | - | 992 | 276 |
| - | - | *C. herbarum* | *A. alternata* | 1739 | 359 |

Airborne fungi, especially in outdoor and hospital environments remain difficult to rapidly detect, despite their potential to contribute to invasive fungal disease and allergic exacerbation in susceptible patient populations.^34, 35^ Therefore, improved methods for rapid detection of airborne fungi are greatly needed in managing allergic responses and invasive fungal disease. The ortholog analysis presented here provides numerous candidate proteins likely accessible to both the immune system and potential diagnostic assays. Our comparative proteomics approach and ortholog comparison also revealed that much overlap exists between *A. fumigatus* and other allergy-inducing fungi, further complicating the development of future diagnostics and highlighting the issue of cross-reactivity between organisms. The dynamic nature of the surface proteome of fungi adds an additional complexity to this challenging problem. Despite these hurdles, we hope that the datasets assembled here will serve as a resource in guiding future endeavors into the biology of the conidial surface and selection of diagnostic targets, particularly in regard to antibody-based detection and chimeric antigen receptor T cell-based therapeutics, which are proving to be more versatile in combatting infectious diseases.^70^

## Conclusions

The surface of fungal conidia is a dynamic meshwork of macromolecules, including a number of important proteins.^8^ In this study, we identified the surface-exposed and secreted proteome of resting conidia from several allergenic fungal pathogens and searched for orthologs between these organisms. We find that the surfaces of germinating fungi are quite variable, which creates unique challenges for the identification of these organisms during allergy and infection and further confirms that these fungi have sophisticated ways of managing diverse environments. In fact, we show that temperature has a great effect on the surface proteome of *A. fumigatus*, a factor which must be considered in the design of future experiments. We were able to significantly add to our collective knowledge of the surface proteome of allergenic fungi in the early stages of germination and now have a resource that will aid in the design of future diagnostics and therapeutics.

## Acknowledgements

We would like to thank Laura Broschat for excellent technical assistance and Antje Häder for her assistance in creating the Δ*cweA* knockout strain. We would also like to thank Vito Valiante for providing an updated gene annotation for *scwA* and Volker Schwartze for thoughtful discussions throughout this project. We thank Amelia Barber for critical reading of this manuscript. MGB was funded by the EXASENS project 13N13861. This work was supported by the Deutsche Forschungsgemeinschaft (DFG)-funded Collaborative Research Center/Transregio 124 FungiNet (projects A1, B5 and Z2) and the DFG-funded ANR project AfuInf. The authors declare no conflicts of interest.

## Supporting Information

*The following supporting information is available free of charge at ACS website http://pubs.acs.org*

**Figure S1:** Production of Δ*cweA* and Δ*scwA* knockouts in *A. fumigatus*. **Figure S2:** Phylogenetic analysis of *A. fumigatus* cell surface proteins. **Figure S3:** Growth characterization of the Δ*cweA* and Δ*scwA* knockouts. **Figure S4:** Trypsin does not induce major changes in fungal permeability.

**Table S1:** Fungal strains used in this study. **Table S2:** Oligonucleotides and plasmids used in this study. **Table S3:** Surface proteome of germinating *A. fumigatus* conidia. **Table S4:** Surface proteome of Δ*scwA* and Δ*cweA* deletion strains. **Table S5:** Time and temperature-dependent surface proteome of *A. fumigatus* conidia. **Table S6:** Surface proteome and secretome of resting and swollen *A. fumigatus* conidia. **Table S7:** Surface proteome and secretome of resting and swollen *P. rubens* conidia. **Table S8:** Surface proteome and secretome of resting and swollen *A. alternata* conidia. **Table S9:** Surface proteome and secretome of resting and swollen *C. herbarum* conidia. **Table S10:** Ortholog analysis performed using InParanoid software package.

---

For TOC Only

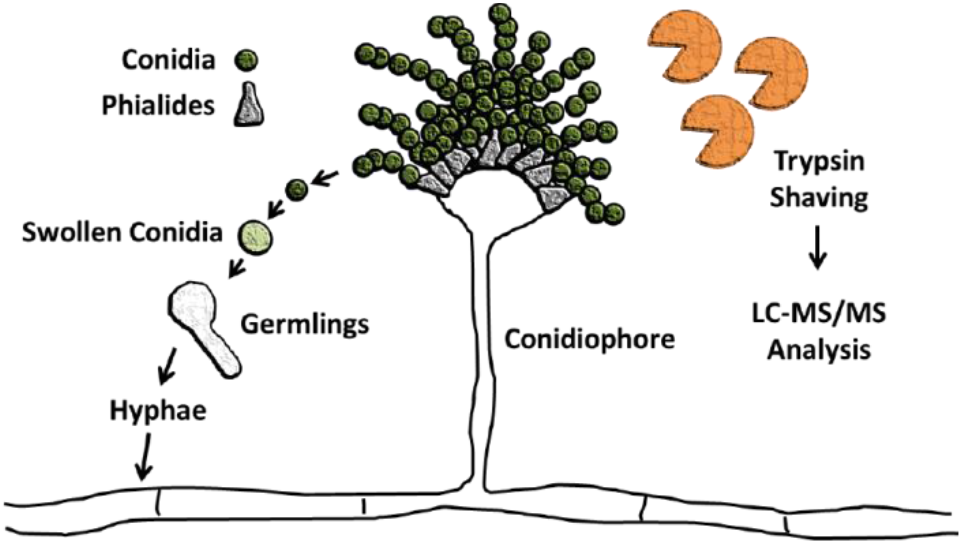

