## Supporting Materials for "The dynamic surface proteomes of allergenic fungal conidia"

*Correspondence:

**Table of Contents for Supporting Information**

*Supporting Figures* *Pages*

**Figure S1:** Production of Δ*cweA* and Δ*scwA* knockouts in *A. fumigatus*. **S-3, S-4**

**Figure S2:** Phylogenetic analysis of *A. fumigatus* cell surface proteins. **S-5, S-6**

**Figure S3:** Growth characterization of the Δ*cweA* and Δ*scwA* knockouts. **S-7**

**Figure S4:** Trypsin does not induce major changes in fungal permeability. **S-8, S-9**

*Supporting Tables*

**Table S1:** Fungal strains used in this study. **S-10**

**Table S2:** Oligonucleotides and plasmids used in this study. **S-11**

**Table S3:** Surface proteome of germinating *A. fumigatus* conidia (Excel File).

**Table S4:** Surface proteome of Δ*scwA* and Δ*cweA* deletion strains (Excel File).

**Table S5:** Time and temperature-dependent surface proteome of *A. fumigatus* conidia (Excel File).

**Table S6:** Surface proteome and secretome of resting and swollen *A. fumigatus* conidia (Excel File).

**Table S7:** Surface proteome and secretome of resting and swollen *P. rubens* conidia (Excel File).

**Table S8:** Surface proteome and secretome of resting and swollen *A. alternata* conidia (Excel File).

**Table S9:** Surface proteome and secretome of resting and swollen *C. herbarum* conidia (Excel File).

**Table S10:** Ortholog analysis performed using InParanoid software package (Excel File).

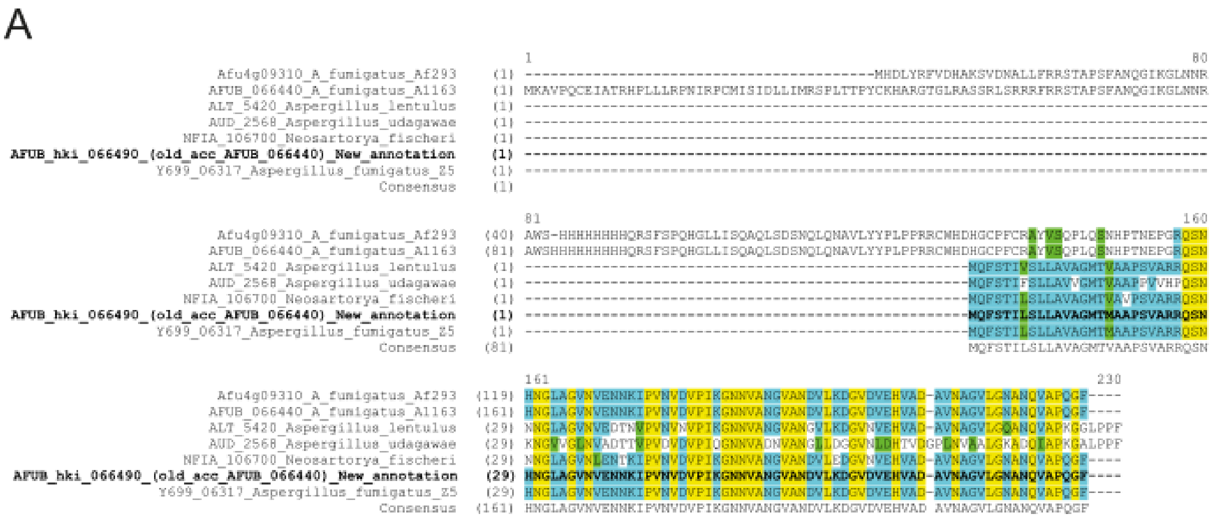

**ScwA Predicted Protein Sequence**

MQFSTILSLLAVAGMTMAAPSVARRQSNHNGLAGVNVENNKIPVNVDVPIKGNNVANGVAND
VLKDGVDVEHVADAVNAGVLGNANQVAPQGF

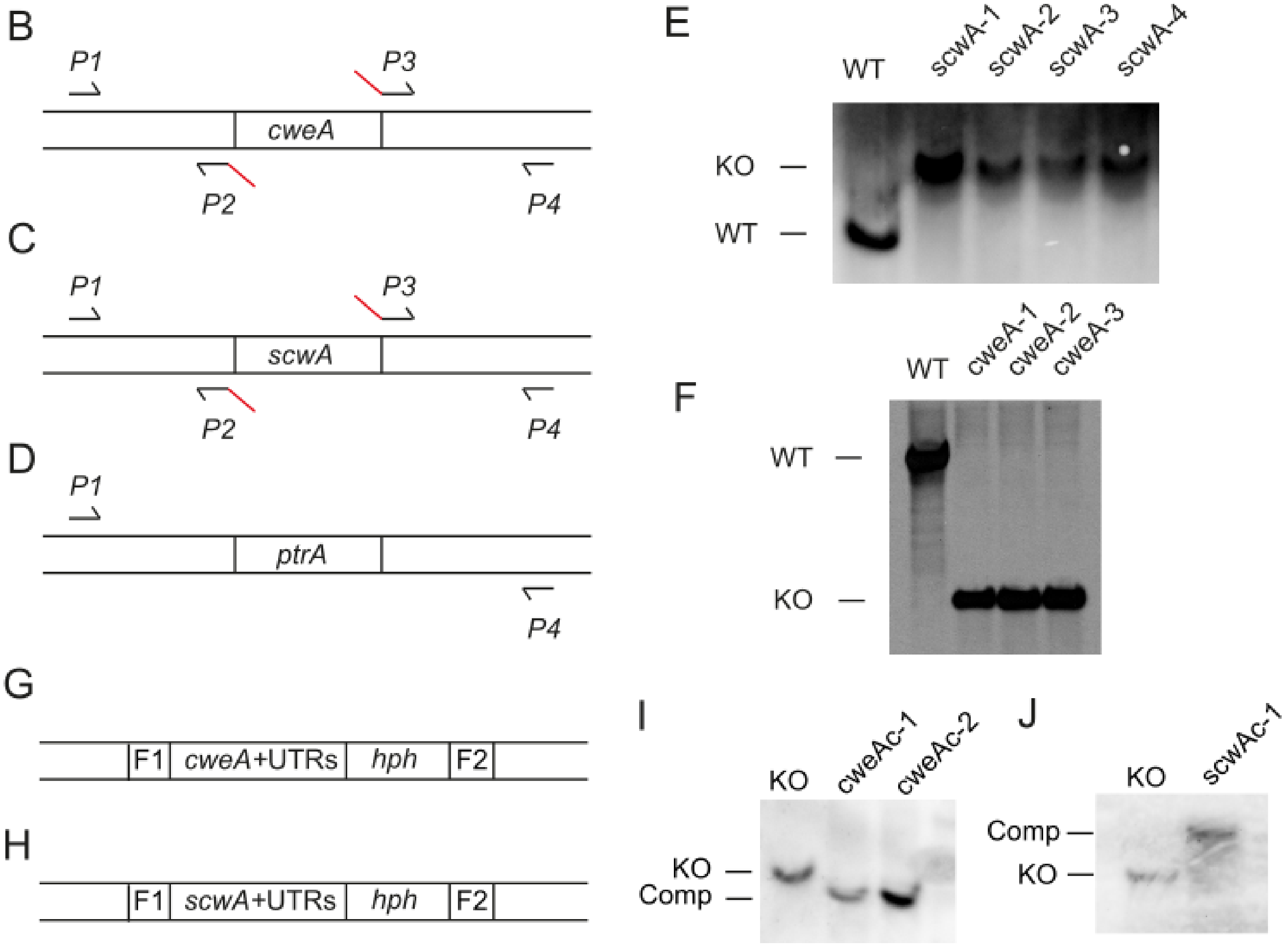

**Figure S1.** Production of Δ*cweA* and Δ*scwA* knockouts in *A. fumigatus*. A) Reannotation of *scwA* (Afu4g09310) from *A. fumigatus* CEA10. For production of genetic deletions of B) *cweA* and C) scwA flanking DNA sequences were amplified using the appropriate primers p1-p4. D) The flanking DNA sequences were then combined with the *ptrA* resistance marker by Phusion polymerase and primers p1/4 to produce a construct with gene flanking regions surrounding the resistance marker. Genetic deletions of E) *scwA* and F) *cweA* were then confirmed by Southern blot analysis after digestion of genomic DNA with *Hin*dIII or *Cla*I, respectively. Complemented strains for G) *cweA* and H) *scwA* were produced by introducing a construct containing a 5' flanking sequence (F1), the gene with both 5' and 3' UTRs, a hygromycin resistance cassette (*hph*), and a 3' flanking sequence (F2) into the original locus of the corresponding knockout strain *via* homologous recombination. Southern blots were performed to confirm complementation of I) *cweA* after digestion of genomic DNA using restriction enzyme *Bam*HI and J) *scwA* using restriction enzyme *Eco*RI.

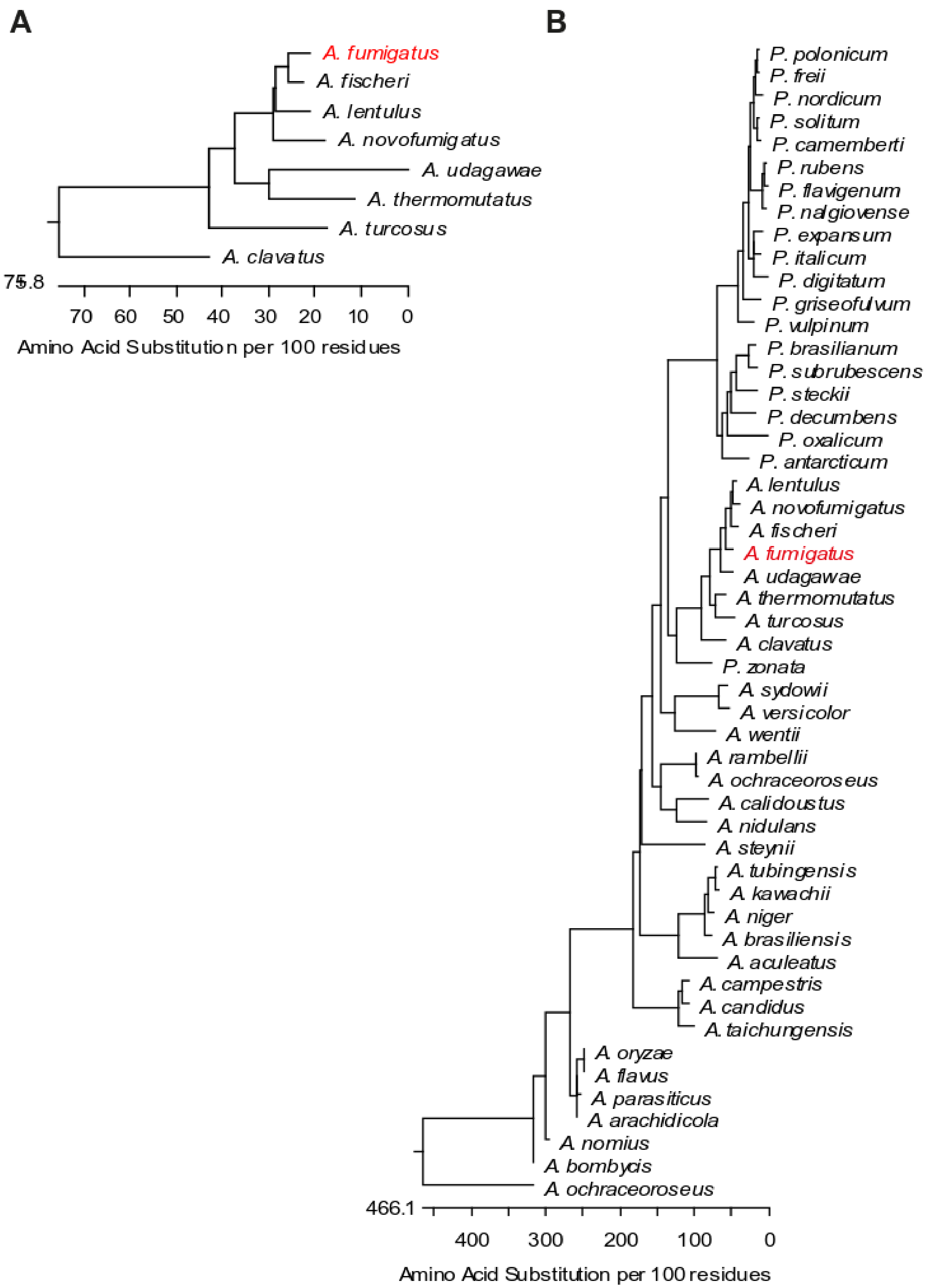
A
A. fumigatus
A. fischeri
A. lentulus
A. novofumigatus
A. udagawae
A. thermomutatus
A. turcosus
A. clavatus
75.8
70 60 50 40 30 20 10 0
Amino Acid Substitution per 100 residues
B
P. polonicum
P. freii
P. nordicum
P. solitum
P. camemberti
P. rubens
P. flavigenum
P. nalgiovense
P. expansum
P. italicum
P. digitatum
P. griseofulvum
P. vulpinum
P. brasilianum
P. subrubescens
P. steckii
P. decumbens
P. oxalicum
P. antarcticum
A. lentulus
A. novofumigatus
A. fischeri
A. fumigatus
A. udagawae
A. thermomutatus
A. turcosus
A. clavatus
P. zonata
A. sydowii
A. versicolor
A. wentii
A. rambellii
A. ochraceoroseus
A. calidoustus
A. nidulans
A. steynii
A. tubingensis
A. kawachii
A. niger
A. brasiliensis
A. aculeatus
A. campestris
A. candidus
A. taichungensis
A. oryzae
A. flavus
A. parasiticus
A. arachidicola
A. nomius
A. bombycis
A. ochraceoroseus
466.1
400 300 200 100 0
Amino Acid Substitution per 100 residues

**Figure S2.** Phylogenetic analysis of *A. fumigatus* cell surface proteins. A) Phylogenetic tree of ScwA sequences produced using the DNAStar MegAlign software with a ClustalW alignment of all sequences deemed significant by NCBI Blastp suite. B) Phylogenetic tree of CweA sequences produced using the DNAStar MegAlign software with a ClustalW alignment of all sequences deemed significant by NCBI Blastp suite.

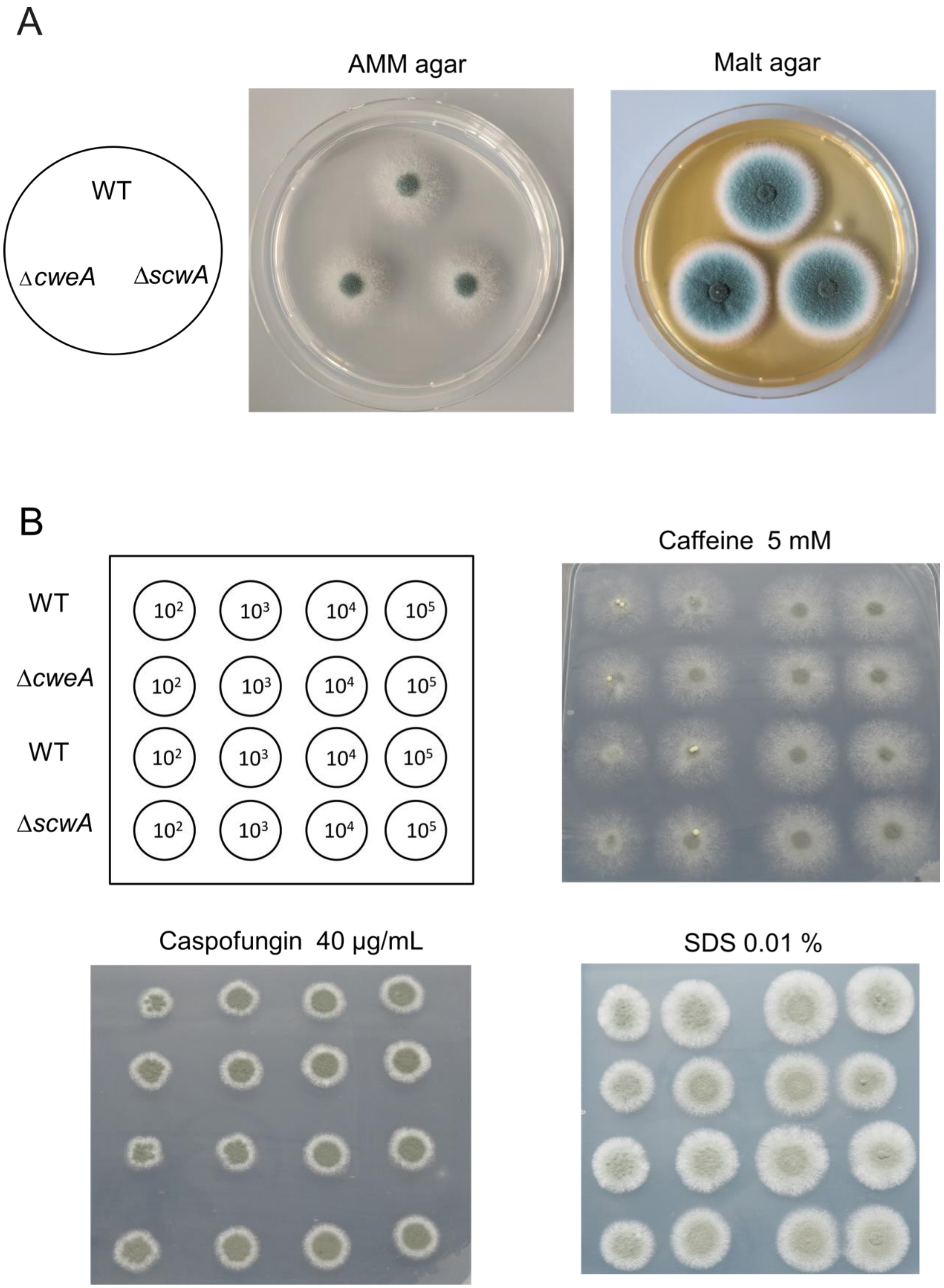

**Figure S3.** Growth characterization of the Δ*cweA* and Δ*scwA* knockouts. A) Growth on AMM and malt agar plates for 48 h. B) Strains were serially diluted and spotted onto AMM agar with the addition of the indicated stressors and grown for 72 h. Experiments are representative data from three separate experiments.

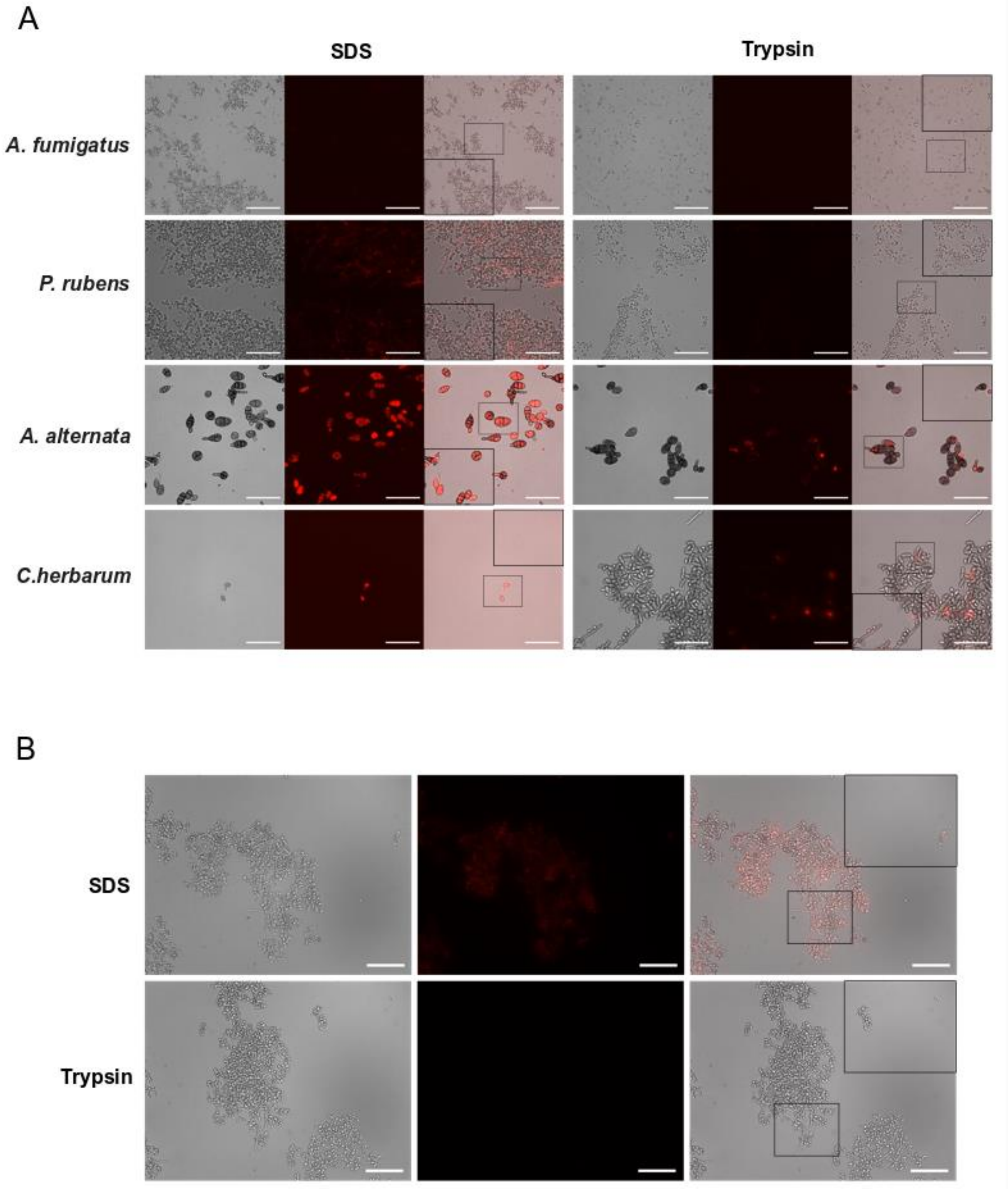

**Figure S4.** Trypsin does not induce major changes in fungal conidial permeability. A) Images show representative fungal conidia stained for 15 min with the cell-impermeable dye propidium iodide (red) following treatment with 1% (w/v) SDS (left) or 5 µg/mL trypsin for 5 min at 37°C (right). B) After 8 hours of incubation, germlings (with hyphal extensions) and swollen conidia also show no lysis after trypsin treatment as performed in A). All scale bars are 50 µm.

**Table S1**. Fungal strains used in this study.

| **Strain** | **Genotype/Phenotype** | **Source** |
|---|---|---|
| CEA10 | *A. fumigatus* wild type | Fungal Genetics Stock Center (A1163) |
| CEA17 | *A. fumigatus* $akuB^{KU80}$ deletion in CEA10 | Da Silva Ferreira et al., 2006 |
| CEA17ΔcweA | Derived from CEA17; *Afu4g09600*::*ptrA*; $PtrA^R$, Δ*Afu4g09600* | This study |
| CEA17ΔcweA + CweA | Δ*cweA* complemented strain | This study |
| CEA17Δ*scwA* | Derived from CEA17; *Afu4g09310*::*ptrA*; $PtrA^R$, Δ*Afu4g09310* | This study |
| CEA17Δ*scwA* + ScwA | Δ*scwA* complemented strain | This study |
| ATCC 66981 | *A. alternata* wild type | American Type Culture Collection |
| ATCC 28089 | *P. rubens* wild type | American Type Culture Collection |
| ATCC MYA-4682 | *C. herbarum* wild type | American Type Culture Collection |

**Table S2**. Oligonucleotides and plasmids used in this study.

| **Oligonucleotide** | **Sequence (5’—3’)** | **Target** |
|---|---|---|
| *Gene Deletion* | | |
| *cweA*-1 | TAACAGAACCTGCCACGACC | *cweA* |
| *cweA*-2 | GGCCTGAGTGGCCATCGAATTCTTCCTCCGACCAGACCCTTC | *cweA* |
| *cweA*-3 | GAGGCCATCTAGGCCATCAAGCGCTGTCATTCGTGCCGTTGA | *cweA* |
| *cweA*-4 | GCAAGCGCAGCAAGATAAGG | *cweA* |
| *scwA*-1 | GCCCACATGCCTCTAGACTC | *scwA* |
| *scwA*-2 | GGCCTGAGTGGCCATCGAATTCCGATCAGCACTCTGGTACTG | *scwA* |
| scwA-3 | GAGGCCATCTAGGCCATCAAGCGGAGATGGCTGGATATAGTC | *scwA* |
| scwA-4 | GCGAGAACCAGAGCGCAAAC | *scwA* |
| *prtA*-1 | GAATTCGATGGCCACTCAGGCC | *ptrA* |
| *prtA*-2 | GCTTGATGGCCTAGATGGCCTC | *ptrA* |
| *Complementation* | | |
| scwA_F | TATCACGAGGCCCTTTCGTCAAGCAAGACCAGAACCTGAAATC | *scwA* |
| scwA_hph_R | GCTCCCCAGGGCTTCTAGTGGATATCTG | *scwA* |
| scwA_hph_F | GCAAAGGAATATGAAGGGCTTTCCCGAAAG | *scwA* |
| scwA_R | CATCGTCACCAACCACACGTCGCGCGTTTCGGTGATGAC | *scwA* |
| hph_scwA_F | GGGCTTCTAGTGGATATCTGCAGAATTCGC | *hph* |
| hph_scwA_R | GTCCGAGGGCAAAGGAATATGAAGGGCT | *hph* |
| pUC19_F | TCGCGCGTTTCGGTGATG | *pUC19* |
| pUC19_R | GTATCACGAGGCCCTTTCGTC | *pUC19* |
| cweA_F | TATCACGAGGCCCTTTCGTCCTAACAGAACCTGCCACG | *cweA* |
| cweA_hph_R | CTAGCTTACGAAGATCAAGAATTTCGCCAGTGTG | *cweA* |
| cweA_hph_F | CAAAGGAATAGTCAGCCATGTGTGGTCTC | *cweA* |
| cweA_R | CATGAGTTTATTAGGGAGCATGAGTCGCGCGTTTCGGTGATGAC | *cweA* |
| hph_cweA_F | TCAAGAATTTCGCCAGTGTGATGGATATC | *hph* |
| hph_cweA_R | GTCCGAGGGCAAAGGAATAGTCAGCCATG | *hph* |
| ***Plasmids*** | ***Source*** | |
| *pUC19* | New England Biolabs | |
| *pUC19_scwAc* | Complementation plasmid produced in this study | |
| *pUC19_cweAc* | Complementation plasmid produced in this study | |
